# Mycobacterial EsxG·EsxH (TB9.8·TB10.4) peptides as a subunit vaccine to booster BCG vaccination in an experimental model of pulmonary Tuberculosis

**DOI:** 10.1101/2024.12.12.628125

**Authors:** Constanza Estefania Martínez-Olivares, Vasti Lozano-Ordaz, Dulce Mata-Espinosa, Jorge Alberto Barrios-Payán, Ángel Elías Ortiz-Cabrera, Yadira Rocio Rodríguez-Miguez, Rogelio Hernández-Pando

**Author notes:** **Corresponding authors**: (CEMO), (RHP).

## Abstract

The attenuated *Mycobacterium bovis* bacillus Calmette-Guérin (BCG) vaccine is currently the only validated vaccine against tuberculosis (TB). In a previous study, we conducted an *in-silico* selection of four peptides (G1, G2, H1, and H2) derived from the mycobacterial protein antigens TB10.9·TB10.4 (EsxG·EsxH). Bioinformatic analysis and molecular dynamic simulations predicted these epitopes could be loaded into a MHC-II complex, inducing T and B cell activation. The present study aimed to experimentally validate these peptides as subunit vaccines by determining their cytotoxicity, immunogenicity, and protective efficacy against *Mycobacterium tuberculosis* (Mtb) in mice when administered as a booster to BCG vaccination. Mice were vaccinated with BCG and, two months later, were subcutaneously immunized with either peptide G1, G2, H1, or H2. One-month post-immunization, mice were challenged with the reference strain H37Rv of moderate virulence or the hypervirulent clinical isolate 09005186. After vaccination and before the challenge, the spleen and lung cells were harvested and stimulated *in vitro* with the corresponding peptide to measure cytokine expression in CD4^+^, and CD8^+^ T cells, as well as the phenotypes of activated effector T cells, proliferative senescence, central and periphery memory CD4^+^ and CD8^+^ cells. Additionally, specific IgG antibody titers elicited by each peptide were measured using ELISA. Compared with animals vaccinated only with BCG, boosting BCG vaccination with these peptides provided enhanced protection by significantly prolonging the mice survival, reducing the bacillary load, and decreasing tissue damage (pneumonia). These findings contribute to the broader understanding of peptide-based subunit vaccines and highlight the potential for tailored approaches to enhance protective immunity.

## Introduction

The attenuated *Mycobacterium bovi*s bacillus Calmette Guerin (BCG) is the only validated vaccine against tuberculosis (TB). BCG was developed in 1921 and is still widely used due to its effectiveness in protecting infants against meningeal and miliary TB. However, its ability to protect adults against prevalent pulmonary TB is limited because BCG induces considerable variability in protection and has limited immunologic memory [1,2]. Thus, the design and testing of new and more efficient vaccines against TB is urgently needed.

Cell-mediated immunity plays a crucial role in coordinating and regulating the immune response to eliminate *Mycobacterium tuberculosis* (Mtb), the causative agent of TB [3]. Significant evidence supports the notion that the polarized response of CD4^+^ cells, specifically the T helper type 1 (Th1) response, is pivotal in protecting against Mtb [4,5]. CD4^+^ T cells activate macrophages, enhancing their ability to defend against microbes. They also attract additional immune cells to sites of infection and inflammation and promote the cytotoxicity of CD8^+^ T cells, which are essential for eliminating infected cells by inducing apoptosis through the action of perforins and granzymes [3]. Considerable discussion exists about the importance of multifunctional T cells, characterized by producing several cytokines instead of just one. Some studies propose that these cells play a crucial role in protecting against TB, while others suggests that they are closely associated with active TB rather than protection [6]. The number of antigen-specific T cells during chronic Mtb infection remains relatively stable, with CD4^+^ T cells proliferating widely. This suggests a high turnover of effector CD4^+^ T cells, indicating that these cells are either depleted or terminally differentiated, and the maintenance of the T cell response depends on the continued replacement of effector T cells [7]. Surface receptors correlate with functional exhaustion and terminal differentiation of effector cells, such as the programmed death receptor 1 (PD-1) and the killer cell lectin-like receptor (KLRG1). PD-1 and KLRG1 identify differentiated, non-depleted CD4^+^ T cells during Mtb infection. KLRG1 expression identifies short-lived, terminally differentiated effector T cells with minimal proliferative capacity, while PD-1 expression identifies activated effector cells with greater proliferative capacity than KLRG1-expressing cells [7]. Meanwhile, humoral immunity is often considered to have lesser protective significance. Nevertheless, substantial recent evidence demonstrated the crucial role of B cells and antibodies in modulating the immune response against TB. B cells acts as antigen-presenting cells to T cells, secrete antibodies, and regulate inflammation through cytokine production [5]. Thus, it seems that a balanced interplay between humoral and cellular immune response is imperative for efficiently controlling and eliminating of Mtb [8].

Indeed, BCG elicits a Th1 response characterized by the secretion of crucial protective cytokines, such as tumor necrosis factor-alpha (TNF-α), gamma interferon (IFN-γ), and interleukin-2 (IL-2), among others. Additionally, there is a moderate expression of cytotoxic mediators like granulysin and perforin produced by CD8^+^ T cells [1]. BCG is also efficient at stimulating innate immunity, with macrophage activation and trained immunity being its most significant activities [9]. Unfortunately, BCG protection has decreased over time and is associated with the predominant activation of effector immune memory over central memory [10]. Due to the limitations of BCG, the pursuit of novel vaccines with improved attributes remains a priority. Among the vaccine categories in development, four types stand out: 1) live attenuated vaccines, 2) whole or fragment inactivated cells vaccines, 3) protein subunit vaccines, and 4) viral vector vaccines. Notably, subunit vaccines are a frontrunner in clinical phases and show high promise, addressing safety concerns and refining antigen selection [5].

However, owing to their use of a narrow range of antigens, subunit vaccines might have a reduced ability to stimulate a broad immune response. To counter this, adjuvants are commonly employed to bolster the immune response they provoke. Typically, subunit vaccines are administered as a booster following the primary BCG vaccination. This approach aims to enhance protection or extend its duration. Subunit vaccines are also being assessed for their therapeutic potential [5].

Immunodominant antigens recognized by T cells, such as ESAT-6, CFP-10, Ag85A, Ag85B, and Tb10.4 [5], have been used to develop subunit vaccines. In a previous study, we conducted an *in-silico* selection of four epitopes derived from the mycobacterial antigens TB10.9·TB10.4 (EsxG·EsxH). Bioinformatic analysis and molecular dynamic simulations predicted that these epitopes could be loaded into a MHC-II complex, inducing T cell activation and, potentially, B cell activation [11]. EsxG·EsxH proteins are pivotal in bacterial iron uptake and adaptation to low zinc environments. Thus, the production of blocking antibodies against these proteins could interfere with mycobacterial metabolism, contributing to bacterial killing. Furthermore, these mycobacterial proteins are known for inhibiting phagosome maturation and delaying T cell activation during infection. Consequently, these epitopes are considered potential candidates for use as subunit vaccines. Thus, this study aimed to experimentally validate the peptides G1, G2, H1, and H2 by determining their cytotoxicity, immunogenicity, and protective efficacy against Mtb in mice. The study used a mild virulent reference strain, H37Rv, and a highly virulent clinical isolate, 09005186, for the challenge infection. These peptides were administered as booster subunit vaccines following primary BCG immunization. Additionally, their vaccine efficacy was compared when administrated alongside aluminum hydroxide as an adjuvant.

## 2. Methods

### 2.1. Peptide synthesis and preparation

The selection and sequence of G1, G2, H1, and H2 peptides were pre-established through *in-silico* analysis [11]. These peptides were synthesized (Genscript Biotech Corporation, New Jersey, USA) via custom peptide synthesis. High-performance liquid chromatography confirmed their purity levels at 92-98%. For experimental use, the peptides were dissolved in a solution of 1% dimethyl sulfoxide (DMSO) (Sigma-Aldrich, Saint Louis, Missouri, USA) and sterile water.

### 2.2. Cytotoxicity assay

The murine alveolar macrophage MH-S cell line (CRL-2019, ATCC, Manassas, VA, USA) was cultured in T-75 flasks containing RPMI 1640 medium (RPP12, Caisson Labs, Smithfield, Utah, USA) supplemented with 10% fetal bovine serum (FBS) (Thermo Fisher Scientific Inc., Waltham, MA, USA) and incubated at 37 °C in a 5% CO_2_ atmosphere until reaching 80% confluence.

For experimentation, 15,000 cells were seeded in each well of 96-well plates. After 24 hours, the peptides (G1, G2, H1, and H2) were added at concentrations ranging from 3.9 to 1000 μg/mL. Cells were then incubated for 48 hours at 37 °C in a 5% CO_2_ atmosphere. Subsequently, the cells were washed with phosphate saline buffer (PBS) at pH 7.4. Following washing, 100 μL of RPMI medium supplemented with 10% FBS, and 20 μL of resazurin sodium salt (Sigma-Aldrich, Saint Louis, Missouri, United States) were added to each well. Cells were further incubated for 24 hours at 37 °C. Resazurin was dissolved in sterile PBS at a concentration of 0.15 mg/mL. Cell viability was assessed using a BioTek Epoch 2 microplate reader (Agilent Technologies, Santa Clara, California, USA) at 600 nm. Untreated cells (without peptide) served as the positive control, representing 100% cell viability, while cells treated with 10% DMSO were the negative control, representing 0% viability. The data were normalized to determine the cell survival rate in response to peptide treatment.

### 2.3. Animals

Pathogen-free BALB/c male mice were sourced and maintained in the animal facility of the Salvador Zubiran National Institute of Medical Sciences and Nutrition (INCMNSZ). Mice were housed under controlled conditions, including a 12:12 hour light/dark cycle, humidity maintained between 50% and 20%, a constant temperature of 23 °C, and provided with *ad libitum* access to food and water. The allocation of mice adhered to the principles of replacement, refinement, and reduction (the 3Rs) for ethical animal research [12]. Animal studies were authorized by the INCMNSZ Animal Experimentation Ethics Committee (approval code: CINVA-PAT-1329-14/17-1), in compliance with the Official Mexican Norm NOM 062-Z00-1999 and the ARRIVE standards [13].

### 2.4. Immunogenicity assay

Six groups of three male BALB/c mice, aged eight weeks, were divided into cohorts receiving either a saline solution (as a control) or a BCG mix containing 4,000 colony-forming units (CFU) of BCG Pasteur and 4,000 CFU of BCG Phipps. Two months post-vaccination, mice were subjected to a subcutaneous injection of 1 μg of either G1, G2, H1, H2, or a mixture of the peptides (mix) at the base of the tail. These peptides, either singly or in combination, were administered once a week for four consecutive weeks. Spleens were harvested on days 7, 14, and 30 post-immunization to assess immunogenicity. Upon collection, spleens were placed in 3 mL of RPMI medium. In comparison, the right lungs were immersed in 2 mL RPMI medium with 1 mg/mL collagenase II (Gibco Life Technologies, Grand Island, New York, USA) and incubated at 37 °C for one hour. Subsequently, spleen and lung tissues were disaggregated using an 18G caliber needle followed by a 21G caliber needle and filtered in a 70 μm strainer (Becton, Dickinson and Company, New Jersey, USA). The resulting cells from both tissues were hemolyzed in a lysis buffer for 5 min at 4 °C, centrifuged, and the pellet was resuspended in 3 mL of RPMI medium. Next, 5×10^6^ cells were stimulated with 5 μg of peptides G1, G2, H1, and H2 for 15 hours at 37 °C, 5 % CO_2_, in the presence of 10 μg/mL of brefeldin A (Thermo Fisher Scientific Inc., Waltham, MA, USA).

After the designated incubation period, 2.5×10^6^ cells were harvested and washed with PBS. For every 1×10^6^ cells, 0.5 μg/mL of purified anti-mouse CD16 / CD32 (Fc Shield) antibody (Tonbo Bioscience, San Diego, California, USA) was added and incubated for 10 minutes on ice. To assess cell viability, 1 μL of the antibody Ghost Dye Violet 450 (Tonbo Bioscience, San Diego, California, USA) was used for every 1×10^6^ cells, and the mixture was then incubated at 4 °C for 20 minutes, shielded from light. Extracellular labeling was carried out using the following antibodies: CD3 AF710 (Tonbo Bioscience, San Diego, California, USA), CD8 BV711 (BD Biosciences, East Rutherford, New Jersey, USA), CD4 BV510 (BD Biosciences, East Rutherford, New Jersey, USA), CD44 FITC (Biolegend, San Diego, California, USA), CD62L APC (Biolegend, San Diego, California, USA), PD-1 PerCP (Biolegend, San Diego, California, USA), KLRG1 PECF594 (BD Biosciences, East Rutherford, New Jersey, USA). The antibodies were diluted at a ratio of 1:20 and incubated for 30 minutes at 4 °C in the dark.

The fixative permeabilizer (eBioscience, San Diego, California, USA) was added to the diluent at a 1:3 concentrator ratio and incubated for 30 min at 4 °C in the darkness. Intracellular labeling was conducted using IFN-γ BV786 (BD Biosciences, East Rutherford, New Jersey, USA), TNF-α BV650 (Biolegend, San Diego, California, USA), and IL-2 PE (BD Biosciences, East Rutherford, New Jersey, USA) antibodies at dilutions 1:100, 1:200, and 1:100, respectively. Incubation occurred at 4 °C in the darkness for 30 minutes. Subsequently, the cells were then resuspended in 500 μL of cold paraformaldehyde, homogenized, and stored at 4 °C. Analysis was performed using a BD LSRII Fortessa cytometer (BD Biosciences, East Rutherford, New Jersey, USA), and we conducted data analysis via FlowJo^TM^ v.10.0.7 Software (BD Life Sciences). For analysis, we selected the lymphocytes zone from FSC-A against SSC-A gate, followed by live cells. The doble-positive cells CD3^+^ CD4^+^ and CD3^+^ CD8^+^ were selected to obtain the percentage of CD4^+^ or CD8^+^ TNF-α^+^, IFN-γ^+^, IL-2^+^, PD1^-^ KLRG1^+^ and PD1^+^ KLRG1^-^.

### 2.5 Animal immunization

Groups of four male BALB/c mice, aged 8 weeks, received vaccinations with BCG mix (4,000 CFU of BCG Pasteur and 4,000 CFU of BCG Phipps). Two months later, mice were subcutaneously immunized with 1 μg, 5 μg, or 10 μg of either synthetic peptide G1, G2, H1, or H2, each administered in a volume of 100 μl. This immunization process was repeated four times for each peptide, with weekly intervals between each administration.

To enhance the peptides’ immunogenicity, another group of animals received vaccinations with the BCG mix, and the peptides were administered in three doses co-administered with 7.5% Inject Alum (Thermo Fisher Scientific Inc., Waltham, MA, USA). The adjuvant was mixed with the peptides for 30 minutes under agitation.

### 2.6 Animal model and challenge

We employed a previously established male model of progressive pulmonary TB [14,15]. The Mtb H37Rv reference strain (ATCC 25618) and clinical isolate 09005186 were cultured in Middlebrook 7H9 broth (BD Biosciences, East Rutherford, New Jersey, USA) supplemented with oleic acid-albumin-dextrose-catalase (OADC) (BD Biosciences, East Rutherford, New Jersey, USA), 0.5% glycerol and 0.5% tyloxapol at 37 °C with gentle agitation (70 rpm) until reaching an optical density (OD) of 1.3 to H37Rv and 09005186, which correspond to the mid-long phase of bacteria measured at 600 nm. At his time, the bacterial cultures were harvested, counted, aliquoted, and stored at −80 °C until use. For the challenge, aliquots of Mtb were thawed and sonicated to prevent clumps formation. Mice were challenged with 250,000 CFU of Mtb H37Rv or clinical isolate 09005186 one month after their last peptide administration. A portion of the inoculum used for the challenge was plated on 7H10 agar plates (BD Biosciences, East Rutherford, New Jersey, USA) supplemented with OADC and glycerol. Twenty-one days later, the CFU counts were performed to confirm the amount and viability of the bacilli. During the challenge procedure, each mouse was anesthetized with 0.1 mL of sevoflurane (Abbott Laboratories, Illinois, USA). A blunt stainless-steel cannula was inserted through the mouth to access the trachea. Intratracheal (IT) administration was confirmed by gently rubbing the tracheal rings with the small ball at the end of the cannula. After being challenged with 250,000 live bacteria, mice were kept upright until they fully recovered. The infection procedure was carried out in a laminar flow hood inside the Animal Biosafety Level 3 (ABSL-3) facilities, and infected mice were housed in groups of four in microisolators.

Four months post-challenge, mice were euthanized via exsanguination under anesthesia, following humane endpoints to prevent animal pain or distress. Mouse survival was monitored throughout the experiment.

### 2.7. Determination of Colony Forming Units (CFU) in Lungs of Tb Mice

The right lungs were utilized to assess the bacillary load by quantifying CFU. Lung tissues were homogenized in 1 mL of PBS-Tween 80 0.05% using a FastPrep tissue homogenizer (MP Biomedicals, Santa Ana, California, USA). Four dilutions of the homogenate were then plated in duplicate on 7H10 agar supplemented with OADC and glycerol. The plates were subsequently incubated at 37 °C in a 5% CO_2_ atmosphere. After twenty-one days, a CFU count was conducted.

### 2.8. Lung area affected by pneumonia

The left lungs were utilized to assess tissue damage corresponding to the area affected by pneumonia. The IT route perfused the right lungs with 100% ethyl alcohol. Parasagittal sections were then dehydrated and embedded in paraffin. Sections of 4 μm thickness were obtained using Leica microtome (Leica Biosystems, Wetzlar, Germany). Subsequently, hematoxylin-eosin staining was conducted. We employed Leica QWin image analysis software (QWin Leica, Milton Keynes, UK) to analyze the histological sections, determining the percentage of lung area affected by pneumonia.

### 2.9. ELISA for antibody titration

ELISA assays were conducted to evaluate antibody production. EIA flat bottom plates (Corning, New York, USA) were coated with 0.5 μg of peptide (G1, G2, H1, or H2) dissolved in 100 μL of 0.1 M sodium bicarbonate, pH 8.0. The coating solution was incubated overnight at 4 °C. Subsequently, the plate was washed three times with PBS-Tween 0.05% (Merck, Darmstadt, Germany). Blocking was performed using a solution of 120 μL of PBS containing 1% of gelatin from cold water fish skin (Sigma-Aldrich, Saint Louis, Missouri, United States) 1 hour, followed by three washes with PBS-Tween 0.05%.

We evaluated a serum pooled from each group. Dilutions of 1:50, 1:500, and 1:5,000 were prepared in 100 μL of PBS–0.1% gelatin, with each dilution placed in duplicate. The serum was incubated overnight at 4 °C. Following incubation, the plate was washed four times with PBS-Tween 0.05%. Subsequently, 100 μL of goat anti-mouse IgG (H+L) HRP conjugate (Bethyl Laboratories, Montgomery, Texas, USA) diluted at 1:5,000 in PBS was added and incubated for 1 hour. After another three times washing with PBS-Tween 0.05%, the ELISA reaction was revealed using OPD Peroxidase Substrate, Sigmafast OPD (Sigma-Aldrich, Saint Louis, Missouri, USA), for 30 min in the dark. The OPD reaction was stopped with 3 M H_2_SO_4_ and read at 492 nm. The antibody titer was calculated using the EC50 of the absorbance obtained in the 1:100 dilution.

### 2.10. Statistical Analysis

The data are presented as the mean ± standard error of the mean (SEM). All data collection was randomized. The normality of data distribution was evaluated using the Shapiro–Wilk normality test, confirming that all data followed a normal distribution. Statistical significance for cytotoxicity, cytokine production, memory, CFU, histology, pneumonia percentage, and antibody titer were assessed using ordinary one-way analysis of variance (ANOVA) with multiple comparisons. A significance level of p < 0.05 was set for all experiments. Survival rates were calculated using the Log-rank (Mantel-Cox) test. Statistical analyses were performed using GraphPad Prism version 8 (GraphPad Software, San Diego, CA, USA).

## 3. Results and Discussion

### 3.1. Cytotoxicity assay

To evaluate the safety and potential toxicity of peptides G1, G2, H1 and H2, we conducted a cellular cytotoxicity assay. Alveolar macrophages were cultured with varying concentrations of the peptides for 48 hours, and cell viability was assessed using resazurin. A cell viability of 100% indicated untreated cells, while 0% represents cells treated with 10% DMSO.

Our results indicate that the four peptides exhibited relatively low cytotoxicity, with noticeable toxicity observed only at high concentrations, especially 1000 μg/mL (Fig 1A-1C). Peptide G1 showed a cell survival rate of 68.9% at 125 μg/mL, decreasing to 44.65% at 1000 μg/mL (Fig 1A). Interestingly, peptide G2 demonstrated the highest cytotoxicity, with a cell survival rate of 69.3% at 15.62 μg/mL, dropping to 39.1% at 1000 μg/mL (Fig 1B). This observation aligns with previous findings where peptide G2 failed to meet *in silico* selection criteria [11]. In contrast, peptides H1 and H2 exhibited lesser cytotoxic effects, with cell survival rates of 50.5% and 49.5%, respectively, at 1000 μg/mL, (Figs 1C and 1D). Thus, peptides derived from the EsxG protein demonstrated varying toxicity levels, with G2 exhibiting the highest. Further comprehensive research is necessary to elucidate these peptides’ toxicity mechanisms fully.

**Fig 1.**
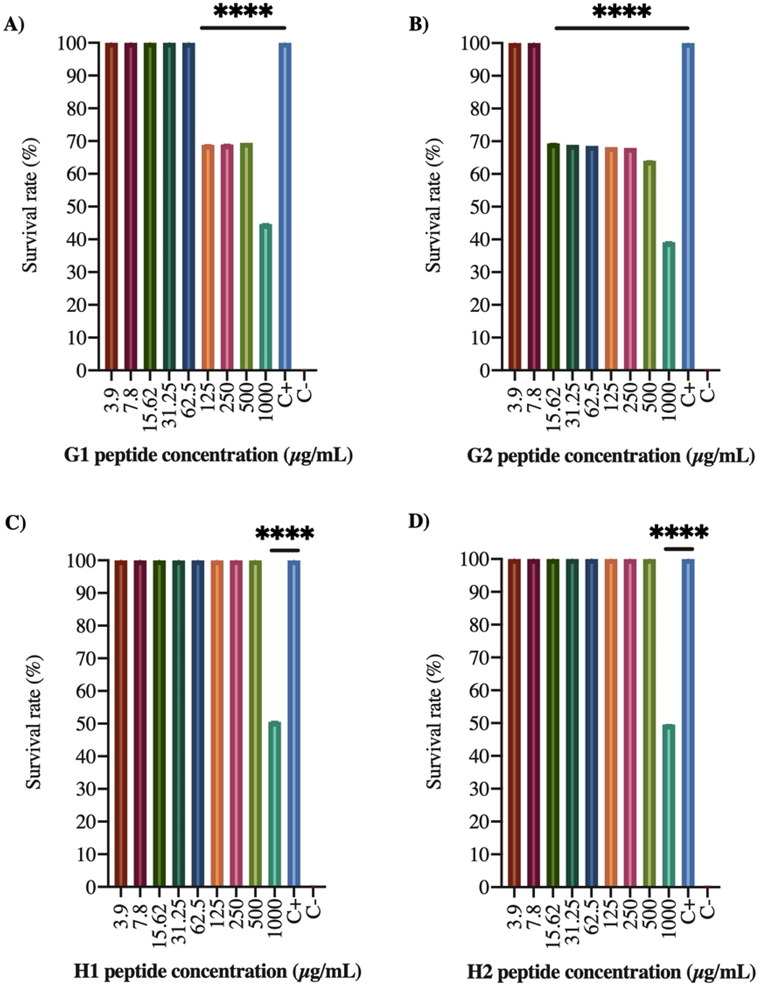
Cytotoxicity assay of peptides G1, G2, H1, and H2. Alveolar macrophages were cultured and incubated for 48 hours with varying concentrations of the indicated peptide. Cell viability was determined using resazurin. Survival data was normalized with 100% viability for untreated cells and 0% viability for cells treated with 10% DMSO. The plots display triplicate data’s average and standard deviation from two independent experiments.

### 3.2. Immunogenicity assay

#### Cytokine production in CD4 and CD8 cells

The immunological response elicited by the peptides was evaluated through subcutaneous immunization with G1, G2, H1, and H2 peptides, either individually or in combination, administered at the base of the tail. Four administrations of 1 µg each were performed at weekly intervals. The dose of 1 µg was chosen based on the peptides’ *in silico* prediction characteristics [11], aiming to use the minimal amount necessary to elicit a specific immune response while avoiding adverse reactions. Spleen and lung cells were harvested and *in vitro* stimulated with the corresponding peptide to identify the CD4^+^ and CD8^+^ T cell populations producing TNF-α, IFN-γ, and IL-2, crucial in the immune response against Mtb. This immune cellular response was evaluated on days seven, fourteen, and thirty post-immunization using flow cytometry. The gating strategy is presented in S1 Fig.

Notably, cytokine production was not detected in CD4^+^ cells of the spleen (Fig 2A). However, immunization with G1 and H2 peptides following BCG vaccination resulted in significantly higher TNF-α production by CD4^+^ cells in the lungs on day 14 post-immunization (Fig 2B). Regarding the cytokine production by CD8^+^ cells of the spleen and lung, no significant differences were observed in the other groups and days evaluated compared to the BCG control group (Figs 2C and 2D). These results suggest that the immunological response triggered by the peptides, whether administered individually or as a mixture, did not consistently induce substantial cytokine production at the measured time points, except in the lungs of groups vaccinated with BCG and boosted with the G1 and H2 peptides on day 14 post-immunization.

**Fig 2.**
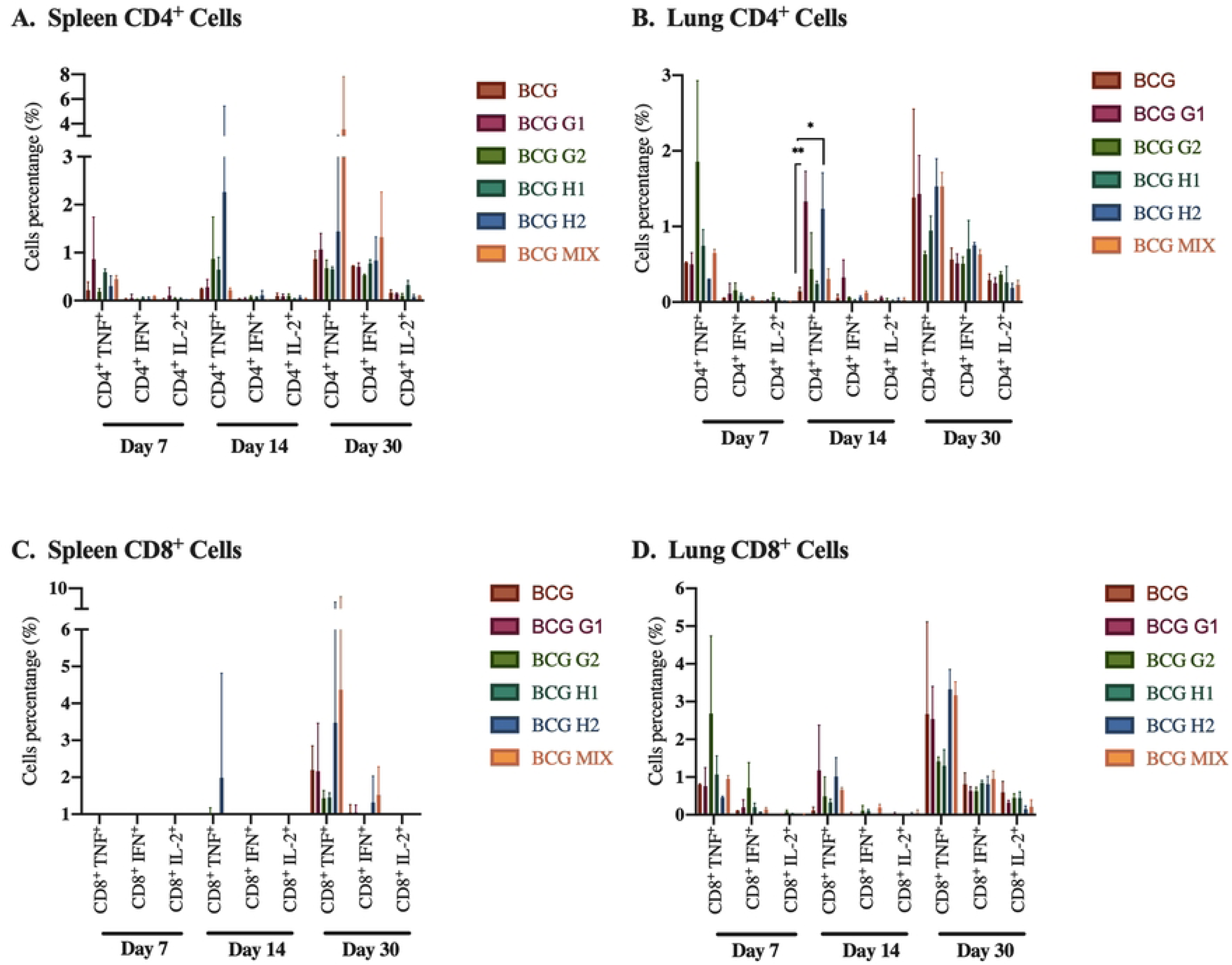
CD4^+^ and CD8^+^ T cells produce TNF-*α*, IFN-*γ*, and IL-2 from the spleen and lung. A) Spleen CD4^+^ cells. B) Lung CD4^+^ cells. C) Spleen CD8^+^ cells. B) Lung CD8^+^ cells. Measurements were taken on days 7, 14, and 30 post-immunization with BCG (control group), or with the indicated peptide or a mixture of peptides.

While our results did not reveal IFN-γ production, it is well established that IFN-γ plays a crucial role in defending against TB [16]. It has become clear that IFN-γ, although essential, is insufficient to ensure protection against Mtb, as evidence in both animal models and humans [16]. The strength and occurrence of the vaccine-triggered IFN-γ response do not reliably forecast immunity vaccines [4]. However, we did observe TNF-α production upon immunization with G1 and H2 peptide in the lungs on day 14, which is significant in immune protection responses against TB. It has been noted that the ablation of TNF-α in lymphoid cells renders protective immunity against TB ineffective [17]. Because IFN-γ and TNF-α have pleiotropic functions in T cell differentiation, cellular trafficking, and macrophage activation and are produced by various immune cells, the mechanisms by which they confer protection are not fully understood [18]. Previous studies have shown that IL-2 inhibits Mtb replication [19]. Recombinant IL-2 therapy in TB patients has demonstrated promising results, leading to clinical improvement and a reduction in bacterial load [20,21]. Enhanced expression of IL-2 in BCG-vaccinated groups enhances the immune response against Mtb infection [19]. However, controversial data suggest that the induction of IL-2 alone may not be sufficient to protect against TB infection [22]. Furthermore, there have been reports of a lack of correlation between the induction of a robust response of Mtb-specific CD8^+^ T cells and protective immunity in experimental vaccines [23,24]. Therefore, cytokine production should not necessarily be synonymous with protection against TB.

There is an ongoing debate about the expression of multifunctional cells’ role in protection. Some studies emphasize their importance in protection against TB, while others associate them with active TB [6]. Our research could not identify multifunctional populations in both CD4^+^ and CD8^+^ cells. It has been demonstrated that the frequency of Th1 cells does not consistently correlate with protection against TB. This lack of a clear protective correlation poses a significant challenge in advancing better vaccines [4].

Our findings on cytokine expression in CD4^+^ and CD8^+^ T cells underscore the immune response’s complex and context-dependent nature. These responses can vary significantly in cytokine production profiles across different tissues and at various time points following immunization. The limited cytokine response observed in *in vitro* stimulation suggests that the peptides might need to be more effective in eliciting a robust immune response in protecting against TB. This data emphasizes the potential impact of the peptide sequence, genetic background, and tissue location on cytokine production, which may have implications for understanding immune responses in TB. Additional research is imperative to draw comprehensive conclusions regarding the role of these peptides in both CD4^+^ and CD8^+^ cell responses and to explore their potential benefits in combating TB. Unraveling the mechanisms driving these observed patterns can offer valuable insights for optimizing immune interventions.

One of the limitations of our study was the inability to evaluate antigenicity using a larger quantity of peptides for immunization. Although prior research has suggested that stimulation may not always depend on concentration, we expected that immunizing with a greater antigen quantity might produce cytokine in other experimental groups.

#### Effector and central memory cells

The success of vaccination is primarily attributed to its ability to elicit memory cells after the initial exposure to an antigen. Therefore, we assessed lymphocyte populations to discern effector memory (CD44^high^ CD62L^-^) and central memory (CD44^high^ CD62L^+^) cells. In CD4^+^ lymphocytes from the spleen, no differences were observed among the groups vaccinated with BCG and boosted with peptides (Fig 3A). However, the group immunized with BCG and then boosted with the G2 peptide exhibited increased lung effector memory cells by day 30 post-immunization (Fig 3B). Regarding CD8^+^ effector memory cells in the spleen, no differences were found between treated and control groups vaccinated with BCG across the three-time points evaluated (Fig 3C). Nevertheless, within the BCG-vaccinated and peptide-boosted cohorts, we observed an increase in this cell population in the subset that received a boost with the H1 peptide by day 30 post-immunization (Fig 3D). These findings underscore the intricate and tissue-specific dynamics in memory cell development in response to various immunization strategies. Understanding these variations in memory cell populations could aid in designing optimized immunization strategies that promote robust and balanced immune responses across different tissues.

**Fig 3.**
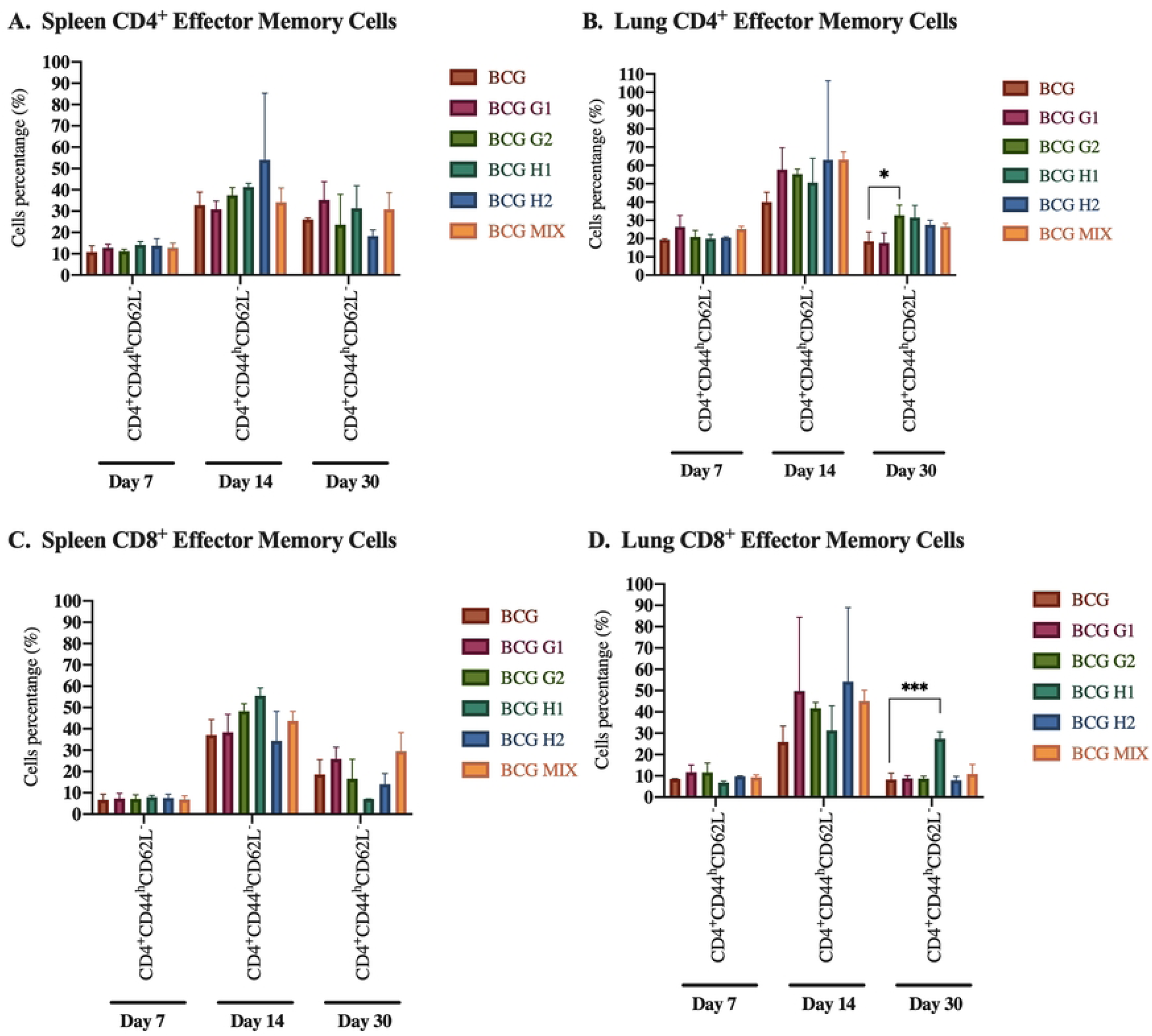
Effector memory CD4^+^ and CD8^+^ in the spleen and lung. A) Spleen CD4^+^ effector memory. B) Lung CD4^+^ effector memory. C) Spleen CD8^+^ effector memory. D) Lung CD8^+^ effector memory. Measures on days 7, 14, and 30 post-immunization with BCG (control group) or BCG plus peptide G1, G2, H1, H2, or a mixture.

Additionally, the evaluation of CD4^+^ central memory cells from the spleen and lung showed no significant differences between the control group and peptide boosters (Figs 4A-4B). This lack of variation suggests that the peptides might not be eliciting distinct effects on these specific immune cell populations. However, an intriguing trend was observed in the spleen’s CD8^+^ central memory cell population. This population decreased when we boosted the BCG-vaccinated groups with the H1 peptide on day 30 post-immunization (Fig 4C). This variability in response indicates that the choice of peptide and timing of immunization can significantly impact the CD8^+^ central memory cell population in the spleen. Conversely, no differences were observed in the CD8^+^ central memory cell population of the lung at the three time points evaluated in groups vaccinated with BCG and boosted with peptides (Fig 4D).

**Fig 4.**
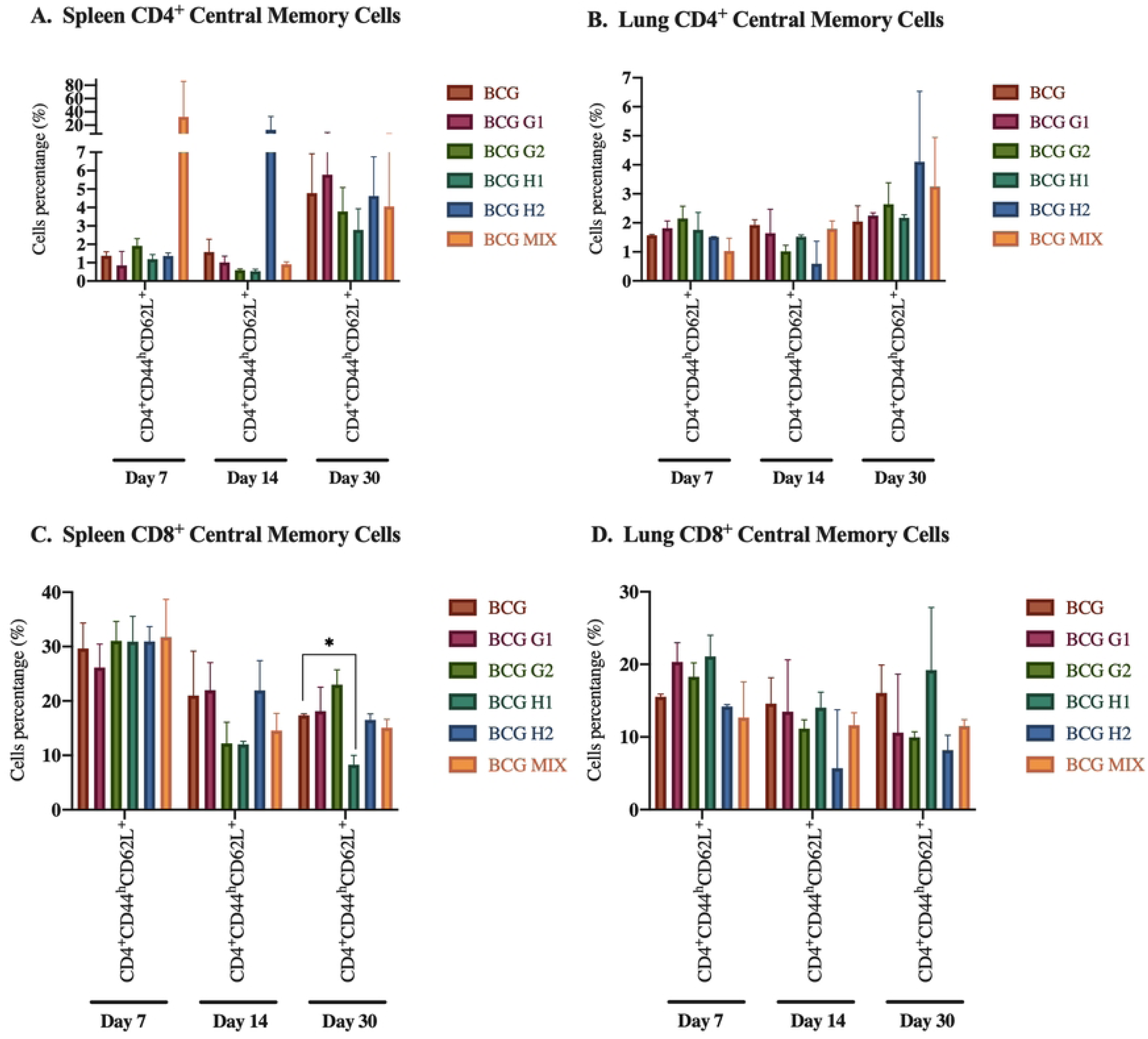
Central memory CD4^+^ and CD8^+^ in the spleen and lung. A) Spleen CD4^+^ central memory. B) Lung CD4^+^ central memory. C) Spleen CD8^+^ central memory. D) Lung CD8^+^ central memory. Measures on days 7, 14, and 30 post-immunization with BCG (control group) or BCG plus peptide G1, G2, H1, H2, or a mixture of them.

Memory responses primarily aim to prevent reinfection or mitigate disease severity [8]. It is worth noting that the BCG vaccine has been reported to predominantly induce effector memory, a characteristic associated with reduced long-term immunity [10]. Given that subunit vaccines excel at generating central memory compared to live vaccines [10], we anticipated an increase in central memory cells. However, we did not observe this boost. Unexpectedly, the group immunized with the H1 peptide exhibited a decline in central memory cells. Previous research has indicated that the development of CD8^+^ memory subtypes can be influenced by factors such as the intensity and duration of antigenic stimulation. Moreover, the durability and maintenance of both effector and central memory cells rely on distinct signaling mechanisms influenced by the specific anatomical regions these cells encounter within the tissues [25]. Understanding these dynamics is crucial for designing effective immunization strategies that elicit a balanced and robust memory response. Regarding with this important point, another significant limitation of our work was the lack of evaluation of these cell subpopulations in peripheral blood samples; perhaps substantial differences could be determined in circulating lymphoid cells, such as central memory CD8^+^ cells that were decreased in the spleen probably because they were circulating in the blood.

#### PD1^+^ KLRG1^-^ and PD1-KLRG1+ in CD4 cells

This study also examined the populations of CD4^+^ PD1^+^ KLRG1^-^ and CD4^+^ PD1^-^ KLRG1^+^ cells in the spleen and lung. In the spleen, we observed an increase in the CD4^+^ PD1^+^ KLRG1^-^ cell population on day 30 post-immunization in the group boosted with the G2 peptide, while we did not observe differences on days 7 and 14 compared to the control group (Fig 5A). No differences were observed in the PD1^-^ KLRG1^+^ population in the spleen (Fig 5B) or in the PD1^+^ KLRG1^-^ population in the lung (Fig 5C). Additionally, the CD4^+^ PD1^-^ KLRG1^+^ cell population showed an increase in the group vaccinated with BCG and boosted with the H1 peptide on day 30 post-immunization, with no differences observed on days 7 and 14 compared to the control groups (Fig 5D).

**Fig 5.**
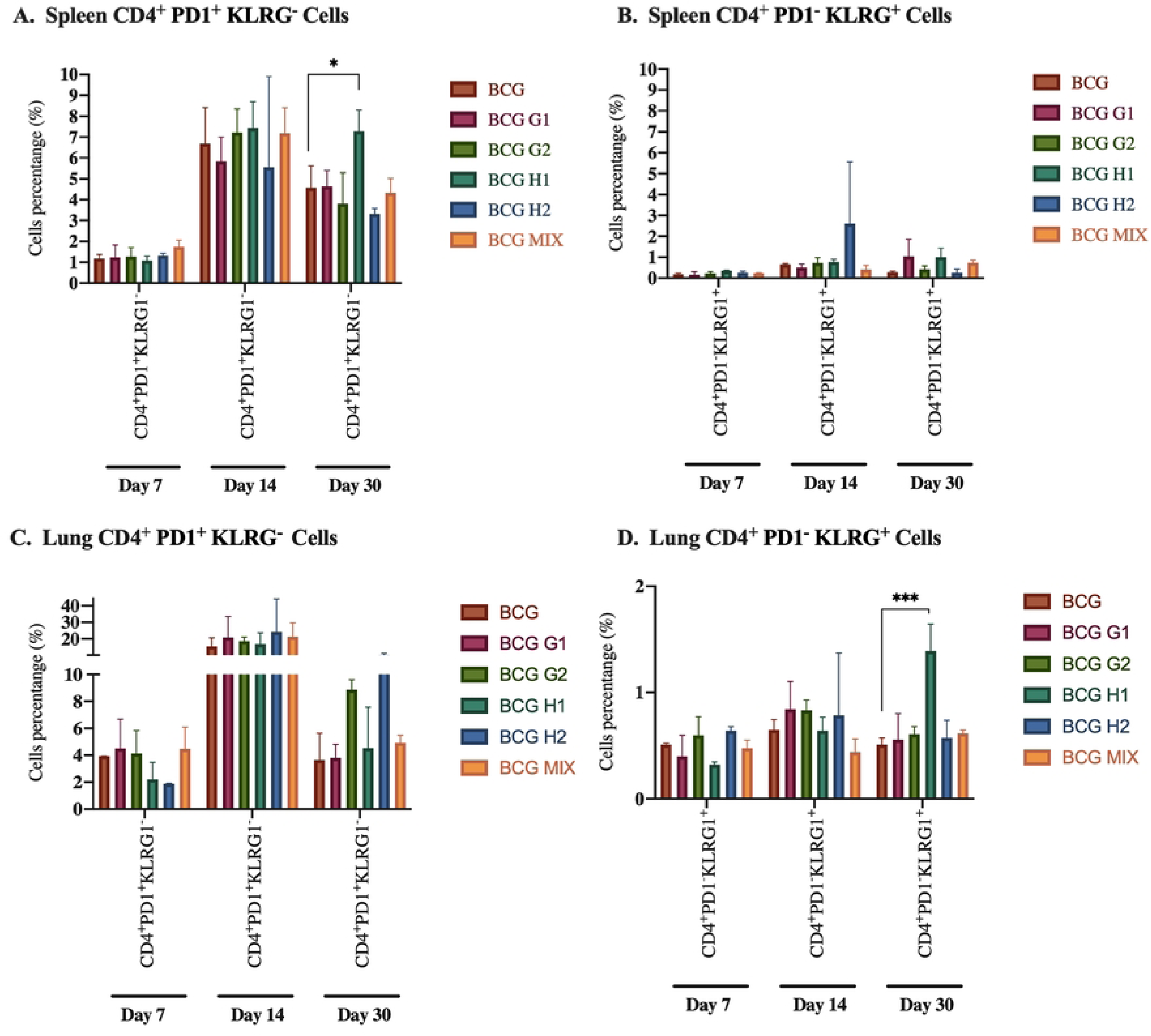
CD4^+^ PD1^+^ KLRG1^-^ and PD1^-^ KLRG1^+^ cells in the spleen and lung. A) Spleen CD4^+^ PD1^+^ KLRG1^-^. B) Spleen CD4^+^ PD1^-^ KLRG1^+^. C) Lung CD4^+^ PD1^+^ KLRG1^-^. B) Lung CD4^+^ PD1^-^ KLRG1^+^. Measures on days 7, 14, and 30 post-immunization with BCG (control group), or BCG plus peptide G1, G2, H1, H2, or a mixture of them.

#### PD1^+^ KLRG1^-^ and PD1^-^ KLRG1^+^ in CD8 cells

Finally, we evaluated the populations of CD8^+^ PD1^+^ KLRG1^-^ and CD8^+^ PD1^-^ KLRG1^+^ cells in the spleen and lung. When mice were vaccinated with BCG and boosted with peptides, no significant differences were observed compared to the control group across the three time points evaluated in the PD1^+^ KLRG1^-^ of the spleen (Fig 6A). In the lung, in the groups vaccinated with BCG and boosted with H1 and H2 peptides, an increase in this specific cell population was observed on day 30 post-immunization (Fig 6C). This observation suggests combining BCG vaccination and peptide boosting impacts the lung’s CD8^+^ PD1^+^ KLRG1^-^ cell population.

**Fig 6.**
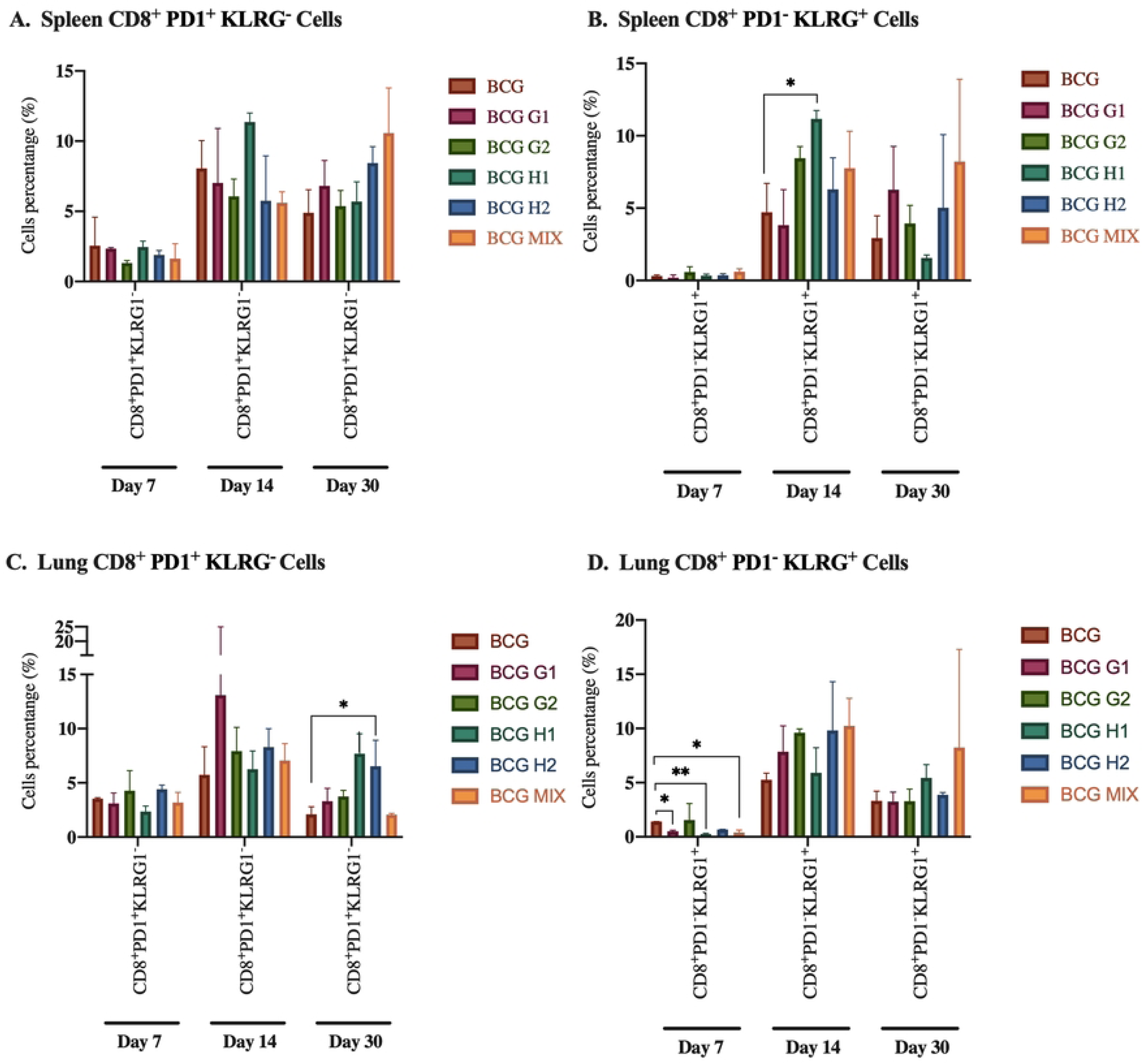
CD8^+^ PD1^+^ KLRG1^-^ and PD1^-^ KLRG1^+^ cells in the spleen and lung. A) Spleen CD8^+^ PD1^+^ KLRG1^-^. B) Spleen CD8^+^ PD1^-^ KLRG1^+^. C) Lung CD8^+^ PD1^+^ KLRG1^-^. B) Lung CD8^+^ PD1^-^ KLRG1^+^. Measures on days 7, 14, and 30 post-immunization with BCG (control group), or BCG plus peptide G1, G2, H1, H2, or a mixture of them.

In the CD8^+^ PD1^-^ KLRG1^+^ cell population, we noted distinctions in the spleen and lung compartments. In the spleen, the group vaccinated with BCG and received a boost with the H1 peptide showed an increase on day 14 post-immunization (Fig 6B). This data could indicate a delayed yet robust response to this vaccination strategy. Interestingly, the cell population decreased on day seven post-immunization in the vaccinated groups that received a boost with the G1, H1, and mixed peptides (Fig 6D). This data suggests a nuanced interplay between the peptides and immune response dynamics within the lung environment.

When employing different vaccination strategies and peptides, the cell populations in the spleen and lung underscore the immune response’s complexity. The correlation between our findings and existing literature is particularly notable. The PD1^+^ KLRG1^-^ phenotype characterizes activated effector T cells [7] and has been linked to better protection against Mtb [4]. Our results show that reinforcing BCG vaccination with the H1 peptide induces this phenotype in CD4^+^ T cells in the spleen. Additionally, in CD8^+^ T cells from the lungs of mice vaccinated with BCG and boosted with the H1 and H2 peptides, this phenotype is observed on day 30 post-immunization. Our data suggest that these vaccination schemes induce a phenotype associated with efficient protection against Mtb.

In contrast, the PD1^-^ KLRG1^+^ phenotype has been associated with proliferative senescence [7] and reduced protective efficacy in TB vaccines [26]. Our results suggest that immunization with the H1 peptide as a boost for BCG generates this phenotype in CD4^+^ T cells in the lung on day 30 post-immunization. It is important to emphasize that studies have reported increased CD8^+^ population after BCG vaccination [27]. In our results, we observed this increase in the spleen cells of the group boosted with the H1 peptide on day 14. Interestingly, in the groups boosted with the G1, H1, and mixed peptides, a decrease in the percentage of this cell population was observed on day 7 post-immunization. These findings underscore the importance of understanding how various immunization strategies can modulate the T-cell response and how this modulation can affect the effectiveness of protection against diseases like TB.

### 3.3. ELISA for antibody titration

Our study comprehensively assesses the immunological responses elicited by peptides across various immunization scenarios. We explored the antibody titers against the G1, G2, H1, and H2 peptides to evaluate their potential as booster candidates for the BCG vaccine. Initially, mice were vaccinated with BCG mix. Two months later, they were subcutaneously immunized with G1, G2, H1, and H2 peptides, individually or in combination, at 1, 5, and 10 μg doses. Immunizations were administered once a week for a month. Additionally, we had groups of mice with peptides combined with aluminum hydroxide. Fifteen days after the final immunization, we assessed the antibody formation against the peptides. Using ELISA, we quantitatively measured specific IgG antibody titers, providing a thorough understanding of the immune responses generated by each peptide and immunization strategy.

In the groups boosted with the G1 peptide, the antibody titer was consistently 1:200, regardless of the presence of an adjuvant (Figs 7A and 7B). In the groups boosted with G2, antibody titers ranged from 1:250 to 1:280 (Fig 7A); when the G2 peptide was administered with aluminum hydroxide, the antibody titers increased slightly, ranging from 1:280 to 1:300 (Fig 7B). For groups immunized with the H1 peptide, with or without aluminum hydroxide, antibody titers ranged from 1:190 to 1:220 (Figs 7A and 7B). In the group vaccinated with BCG and boosted with the H2 peptide, antibody titer ranged from 1:190 to 1:220 (Fig 7A). When aluminum hydroxide was added, the titers slightly increased from 1:220 to 1:250 (Fig 7B).

**Fig 7.**
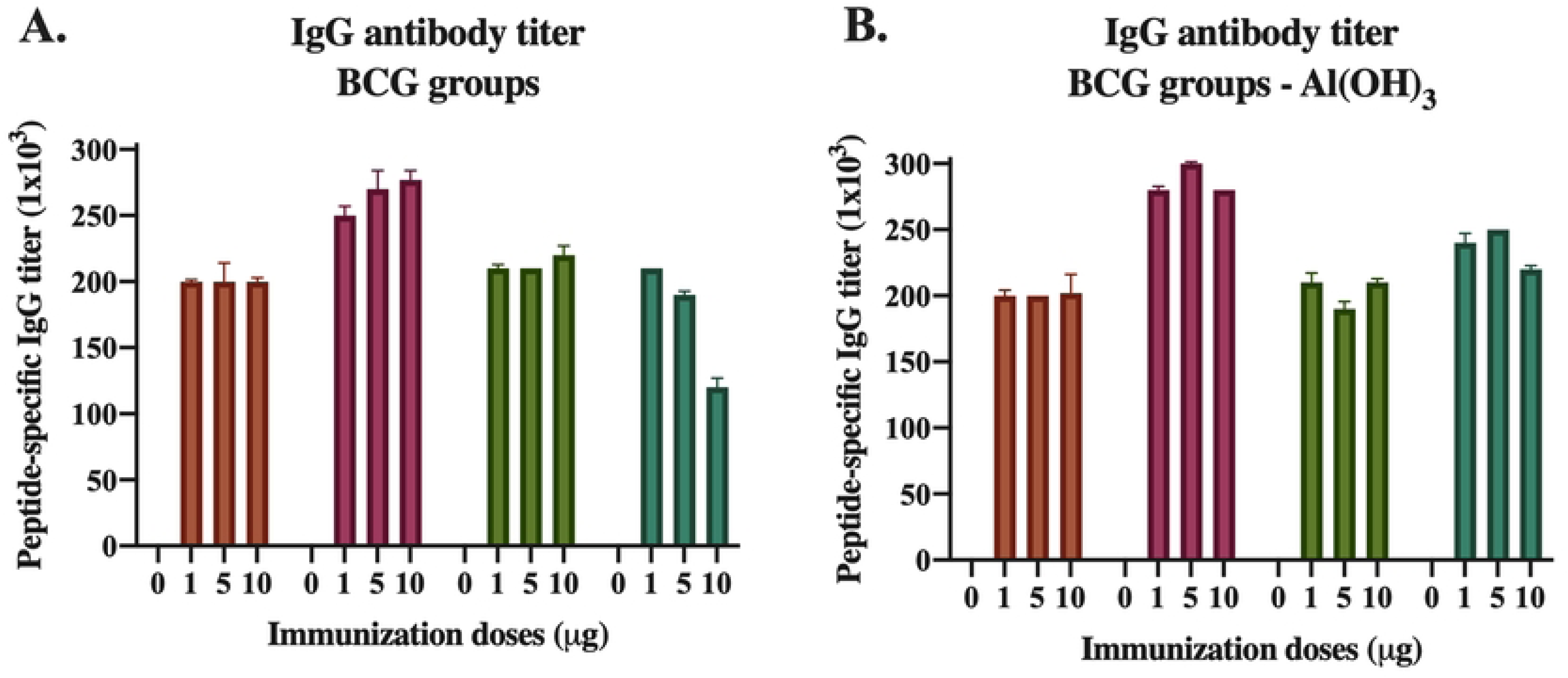
Peptide-specific IgG antibody titer. The x-axis represents the dose used in the immunization (1, 5, or 10 μg) with the G1, G2, H1, or H2 peptides. A pooled serum sample for each group was evaluated. The y-axis represents the antibody titer, calculated using the EC50 of the 1:100 dilution.

In most currently used vaccines, protection correlates with the level of antibodies induced by the vaccine. However, for the BCG vaccine, an increase in antibody titers has not been shown to correlate with protection. Previous reports have indicated that BCG vaccination induces high levels of antigen-specific IgG and long-lasting memory cells, suggesting these antibodies might enhance the opsonization and phagocytosis of Mtb [28].

Our results demonstrated that immunization with BCG followed by a booster with the four evaluated peptides generates a significant IgG antibody response. The G1 and H1 peptides possess inherent immunogenic properties that evoke a stable antibody response unaffected by the immunization dosage or an adjuvant. For the G2 and H2 peptides, aluminum hydroxide slightly enhances the antibody titer, underscoring its capability to augment antibody production. This effect is likely due to the adjuvant creating an environment conducive to a more potent immune response.

It is important to emphasize that this study marks the first instance of evaluating the peptides *in vivo* assays. Previous research on peptides from the EsxG and EsxH proteins primarily focused on assessing cellular response. The notable exception is the study by Coler and colleagues [29], which examined IgG1 and IgG2 antibody titers following immunization with Mtb9.8 (EsxG) formulated with adjuvants AS02A and AS01B. Our findings deepen our understanding of peptide immunology and hold significant potential for optimizing vaccine formulations and strategies. This advancement is crucial for developing effective peptide-based vaccines targeting various diseases. Further investigations are warranted to elucidate the mechanisms driving these diverse immune responses and to guide the rational design of peptide vaccines with enhanced efficacy.

Another limitation of our study was the inability to assess IgG subclasses and the production of isotypes such as IgM and IgA. However, due to the vaccination route employed and the serum assessment, most antibodies are expected to correspond to the IgG isotype. Characterizing the isotypes would be highly useful and essential, as it has been reported that IgG1 promotes bacterial growth, while IgA isotype induces protection [10]. Therefore, focusing on humoral immunity could provide new immunological insights to guide vaccination strategies.

### 3.4. Vaccinated mice response to *Mycobacterium tuberculosis* challenge

Expanding our investigation, we evaluated the protective efficacy of the peptides against bacterial infection. One month after the last immunization, we conducted a challenge using 250,000 CFUs of Mtb H37Rv. Mice survival was monitored for four months following the challenge. No deaths were observed in the group of mice that received BCG vaccination or the peptide boost, nor in the group boosted with peptides and aluminum hydroxide (S2 Fig).

Mice were immunized with 1 μg of each peptide or a mixture of them and subsequently challenged with the strain 09005186, a member of the Latin American Mediterranean genotype. This strain is a clinical isolate characterized by its high transmissibility and prevalence in a community in southern Mexico [30,31]. When mice were vaccinated with BCG and then boosted with the peptides, the group boosted with the H1 peptide demonstrated a significant extension in survival by over 13 days. In the BCG-vaccinated control group, the last event occurred on day 67, while in the immunized group, events were recorded until day 70. The last individual in this group was euthanized on day 80 (S2 Fig). These results strongly indicate that the immunization scheme, when combined with the H1 peptide as a booster following BCG vaccination, provides enhanced protection by significantly prolonging the survival of the mice. A limitation of our investigation was the use of higher doses and the lack of evaluation of the bacterial load because the animals succumbed at different times during the experiment.

The results obtained with the 09005186-strain differed from those obtained with the H37Rv strain due to the higher virulence of the former. Intriguingly, when the H1 peptide was administered as a booster following BCG vaccination, it extended the survival of the challenged mice. Recently, a polymorphism has been reported in the A71S residue of *M. Africanum*, corresponding to the EsxH protein and located at the intermolecular interface with the EsxG protein. This region is adjacent to Met72, which, along with Met18, forms a cleft. This polymorphism disrupts the hydrophobic interaction by replacing it with Ser, a polar residue. This destabilization could interfere with the zinc-binding site and lead to alterations in MHC molecule binding, thus impacting protein functionality [32]. Additionally, another study analyzing clinical isolates reported that 86 of the isolates exhibited polymorphisms in the *esxH* gene. The A10T polymorphism in the EsxH protein alters antigen processing and presentation to MHC I, resulting in CD8 cells being unable to recognize infected macrophages and, consequently, unable to provide optimal protection against Mtb [33]. These polymorphisms represent a strategy employed by Mtb to evade the immune response. The effects observed after the administration of the peptides in the employed strains could be a consequence of these polymorphisms. However, to our knowledge, no polymorphic regions have been reported for the *esxG* gene. Further studies are required to delve deeper into this matter, and the impact of including highly polymorphic regions in vaccines is a topic that needs further exploration.

Four months after the challenge with H37Rv strain, mice were euthanized, and we determined the number of CFUs in the right lungs. Peptides were administered as a booster to the BCG immunization to investigate if they could enhance the immune response and improve protection, which was significantly produced depending of the peptide and its dose. Compared to the BCG-vaccinated control group, the group booster with peptide G1 demonstrated a reduction in the bacillary load of 88.70 %, 88.04 %, and 96.74 % at doses of 1, 5, and 10 μg, respectively (Fig 8A). Peptide G2, as a booster, resulted in a reduction in the bacillary load of 67.39 % and 91.30 % at 5 and 10 μg doses, respectively. The 1 μg dose showed no statistical difference compared to the BCG control (Fig 8B). No differences were observed between the groups immunized with 1 and 10 μg of the H1 peptide and the BCG-vaccinated control group. However, in the group vaccinated with 5 μg, the CFU decreased by 50% compared to the control group (Fig 8C). Similarly, there was no difference between the BCG control group and the group boosted with 1 μg of peptide H2. The 5 and 10 μg doses reduced the bacillary load by 86.30%, and 82.26%, respectively (Fig 8D).

**Fig 8.**
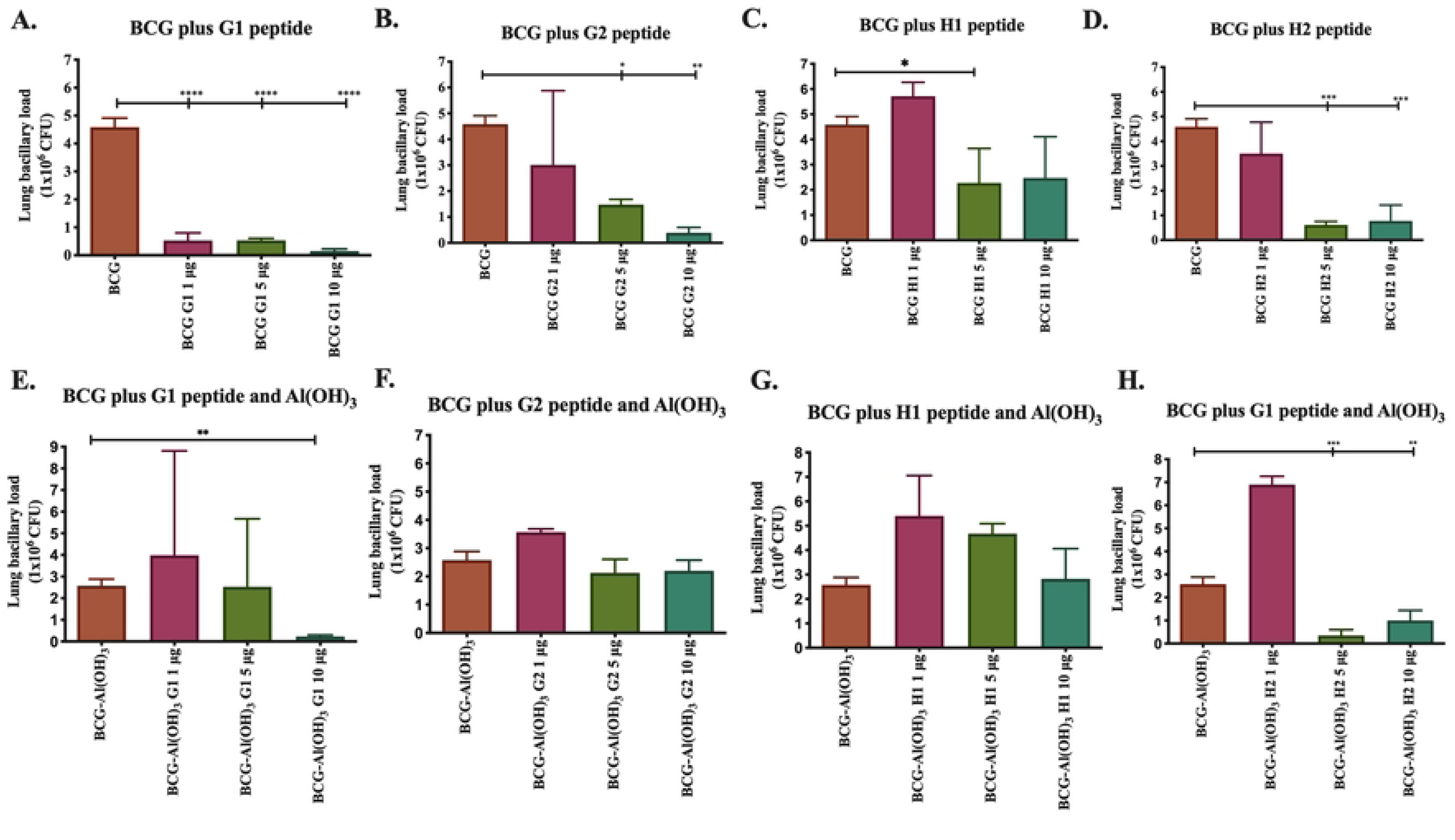
Bacillary load in mice lungs four months after booster immunization with the indicated peptide with or without aluminum hydroxide. Colony-forming units per lung is shown on the y-axis.

When vaccinating with BCG and administering the peptides as aluminum hydroxide boosters, the CFU decrease differed from the results obtained without this adjuvant. Immunization with 10 μg of G1 peptide produced a 91.15% reduction in CFU, while no decrease in the bacillary load was observed in the 1 μg and 5 μg doses (Fig 8E). For the G2 and H1 peptides, there were no differences compared to the control group at any of the three doses administered (Figs 8F and 8G). However, in the group boosted with the H2 peptide, the 5 μg and 10 μg doses resulted in decreases of 86.54% and 57.69%, respectively (Fig 8H).

Finally, histological sections of the left lungs of the mice were prepared to assess the extent of pneumonia. We observed that in the groups receiving BCG vaccinations along with peptide boosters, the pneumonia rates remained similar to the control group that only received BCG vaccinations, except for the groups administered 1 μg of G1, 1 μg of G2, and 1 and 5 μg of H2. Despite some variations, the prevalence of pneumonia remained around 30% (Figs 9A – 9D). In the groups where an adjuvant was used, there was no significant reduction in pneumonia compared to the control group (Figs 9E – 9F).

**Fig 9.**
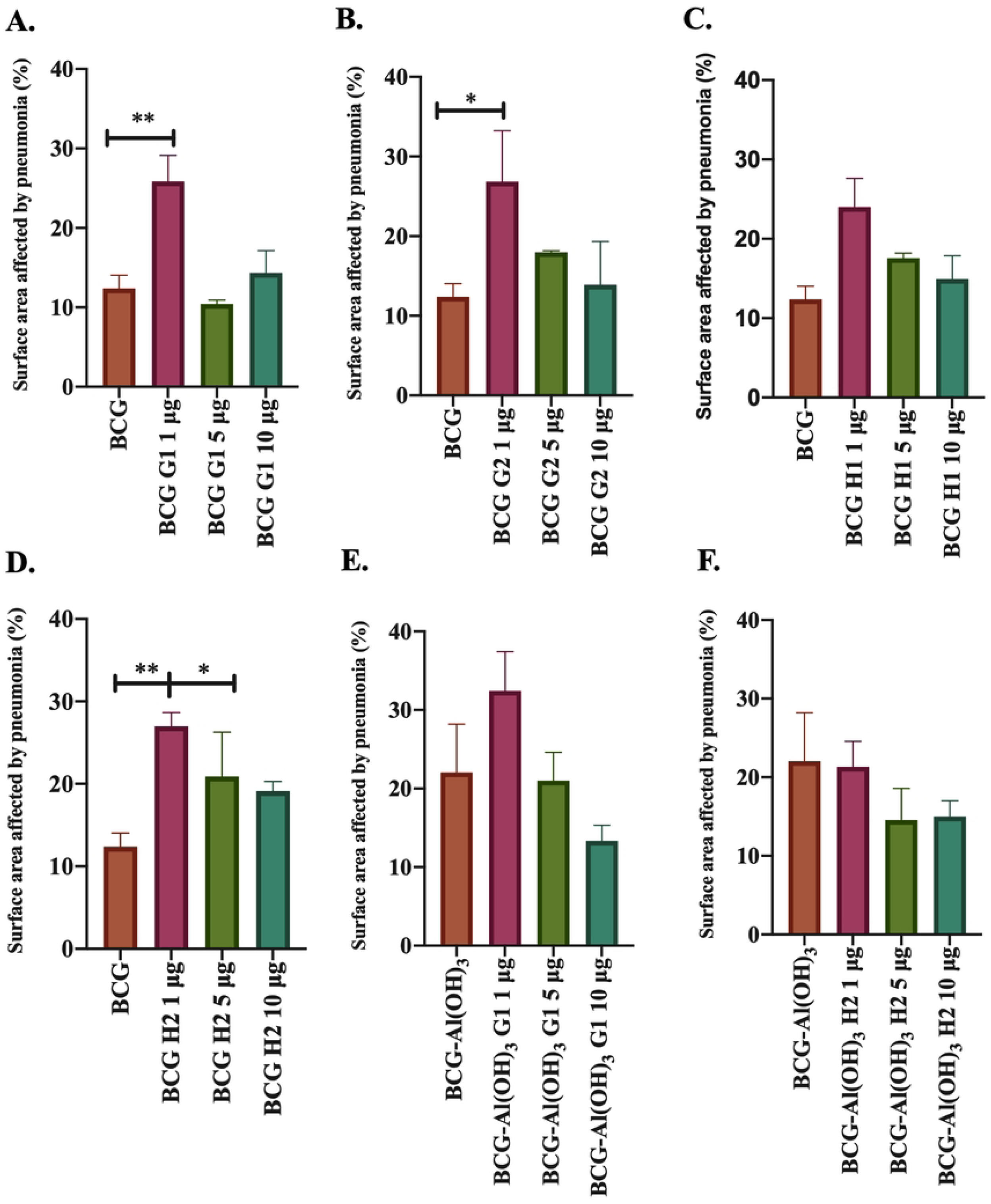
Percent pneumonia in mice’s lungs four months after booster immunization with the indicated peptide with or without aluminum hydroxide. Percent pneumonia per lung is shown on the y-axis.

Subunit vaccines often lack the necessary molecular signals to activate the innate immune system efficiently, resulting in a limited induction of robust adaptive immunity [10]. Our results indicate that the administration of the selected peptides provides protection against Mtb in immunized mice, contrasting with previous findings. In earlier studies, mice were immunized with the IMYNIPAM peptide along with the CAF05 adjuvant, administered intraperitoneally three times at two-week intervals, and challenged with Mtb Erdman seven weeks later did not exhibit a decrease in CFU in the lungs [34]. Conversely, another study demonstrated that intramuscular administration of the EsxG with the AS01B or AS02A adjuvant reduced the bacterial load in the lungs and spleen of C57BL/6 mice challenged with Mtb Erdman. Furthermore, when the protein was administered with the AS02A adjuvant, it extended the survival of guinea pigs [29].

Common adjuvants like aluminum are co-administered to address the peptide limitation and enhance specific immune responses. It is worth noting that aluminum, the only globally approved adjuvant for human use, primarily supports Th2-dependent immune responses but may impede cytotoxic responses and potentially lead to side effects [10,35]. Our results indicate that the co-administration of peptide G1 with aluminum hydroxide enhances protection in terms of bacterial load, as does the 5 μg dose of H2. However, for other peptides, the anticipated synergy was not observed. In the scheme where peptides were used as boosters for BCG vaccination, we noticed that aluminum hydroxide did not significantly reduce bacterial load in most cases.

In general, there appears to be a relationship between the decrease in bacterial load and histological damage, although there are some exceptions. Vaccines against TB that include Th2 components like aluminum hydroxide could worsen the disease or increase susceptibility to TB (Lindblad et al., 1997). Furthermore, it has been demonstrated that administering the subunit vaccine Ag85B-ESAT6 with aluminum hydroxide results in a Th2 response, which does not protect Mtb. In contrast, an adjuvant that induces a Th1 response with the same vaccine elicits a protective Th1-type immune response [36]. These findings suggest that including adjuvants in formulating subunit vaccines against TB should be carefully evaluated [37].

## 5. Conclusion

In conclusion, our study provides comprehensive insights into the cellular response induced by four peptides: G1, G2, H1, and H2, previously selected *in silico* [11]. We observed that the cellular response induced by these peptides, regarding cytokine production and memory, is similar to what has been reported in the literature. Furthermore, we found a significant output of IgG class antibodies when evaluating the humoral response.

As it has been established that peptides alone are incapable of generating a protective immune response, they were administered alongside aluminum hydroxide. Interestingly, we observed that, in most cases, immunization with the adjuvant did not enhance protection against Mtb. However, when these peptides were administered without the adjuvant as a booster for BCG, they significantly reduced the bacillary load in a dose-dependent manner compared to animals vaccinated only with BCG. Despite this improvement, no differences were observed in the extent of pneumonia.

Notably, we observed an inverse protective effect in the groups immunized with G2, H1, and the mixture of peptides when the mice were challenged with the clinical isolate 09005186. However, when the H1 peptide was administered as a booster after BCG vaccination, it extended the survival of the mice, a result that could be attributed to polymorphisms and warrants further investigation.

Our results demonstrate the potential use of peptides G1, G2, H1, and H2 as subunit vaccines against Mtb, suggesting that achieving optimal pathogen control requires the activation of both the humoral and cellular immune responses. To the best of our knowledge, this is the first study to show a protective effect against Mtb using peptides from the EsxG and EsxH proteins. The dynamic nature of immune responses, the intricate interplay between peptides and adjuvants, and the complexities of boosting strategies emphasize the multifaceted considerations in vaccine development. These findings contribute to the broader understanding of peptide-based vaccines and highlight the potential for tailored approaches to enhance protective immunity. Further exploration of the mechanisms behind these responses will be invaluable in guiding the optimization of peptide-based vaccine strategies.

---

**S1 Fig.** Cell population selection strategy for flow cytometry assay.

**S2 Fig. The survival rate of mice immunized with peptides G1, G2, H1, and H2 challenged with H37Rv or 09005186 strains. A.** Groups vaccinated with BCG and boosted with or without 1, 5, and 10 μg of the peptides G1, G2, H1, and H2, challenged with the H37Rv strain. **B**. Groups vaccinated with BCG and boosted with or without 1, 5, and 10 μg of the peptides G1, G2, H1, and H2, in combination with aluminum hydroxide, challenged with the H37Rv strain. **C.** Groups vaccinated with BCG and boosted with or without 1 μg of the peptides G1, G2, H1, and H2, in combination with aluminum hydroxide, challenged with the 09005186 strain.

